# Panel Informativity Optimizer (PIO): an R package to improve cancer NGS panel informativity

**DOI:** 10.1101/2021.06.23.449561

**Authors:** Vincent Alcazer, Pierre Sujobert

## Abstract

Mutation detection by next generation sequencing (NGS) is routinely used for cancer diagnosis. Selecting an optimal set of genes for a given cancer is not trivial as it has to optimize informativity (*i.e*. the number of patients with at least one mutation in the panel), while minimizing panel length in order to reduce sequencing costs and increase sensitivity. We propose herein Panel Informativity Optimizer (PIO), an open-source software developed as an R package with a user-friendly graphical interface to help optimize cancer NGS panel informativity. Using patient-level mutational data from either private datasets or preloaded dataset of 91 independent cohort from 31 different cancer type, PIO selects an optimal set of genomic intervals to maximize informativity and panel size in a given cancer type. Different options are offered such as the definition of genomic intervals at the gene or exon level, and the use of optimization strategy at the patient or patient per kilobase level. PIO can also propose an optimal set of genomic intervals to increase informativity of custom panels. A panel tester function is also available for panel benchmarking. Using public databases, as well as data from real-life settings, we demonstrate that PIO allows panel size reduction of up to 1000kb, and accurately predicts the performance of custom or commercial panels. PIO is available online at https://vincentalcazer.shinyapps.io/Panel_informativity_optimizer/ or can be set on a locale machine from https://github.com/VincentAlcazer/PIO.

## INTRODUCTION

In the era of personalized medicine, molecular analysis of cancer genomes by Next Generation Sequencing (NGS) has become an essential step to establish a precise diagnosis, decipher patient prognosis, and identify therapeutic targets. Although comprehensive analysis of the whole genome has been recently proposed in the routine setting, this approach suffers from many limitations such as lack of sensitivity to detect subclonal mutations and excessive cost [1]. As an alternative, the use of target enrichment to analyze a panel of genes of interest has been widely adopted. However, panel design is a complex task that has to reach an optimal cut-off between multiple constraints. On the one hand, limiting the size of the panel is important to reduce the cost of analysis, and increase the depth of sequencing (*i.e*. increase sensitivity of detection of low variant allele frequency events). This constraint is even more important when working on circulating tumor DNA (ctDNA) because the tumor DNA is diluted in circulating DNA from normal cells and because very high sensitivity is needed to monitor residual disease. On the other hand, a large panel is required to ensure detection of mutations with clinical impact, and optimal informativity (at least one detectable mutation in all patients). Whereas the list of genes with clinical impact in a given cancer is well described and consensual [2], there is no generic tool to optimize the informativity of the panel.

We present herein the Panel Informativity Optimizer (PIO), an open-source, publicly available software to optimize panel informativity. Based on individual mutational profiles available in public cancer genomics databases or in private databases, PIO enables the design of an optimal panel (defined at the level of whole genes or exons) to achieve absolute informativity (100% of the patients with at least one mutation). In addition, PIO can be used to assess the informativity of a given panel in a cancer of interest. PIO also includes everal other features, such as the exploration of mutation frequency and co-occurrence. By combining metrics on informativity and panel length, PIO can help make accurate choices in the design of panels or to benchmark commercial solutions. We provide 3 different examples of applications of PIO in real-life settings: size optimization of a 293-gene (802-exon) panel used in Diffuse Large B Cell Lymphomas (DLBCL) [3], benchmarking of five different commercial panels proposed for myeloid malignancies [4], and validation of PIO predictions for a custom 110-gene panel in a local cohort of 714 Acute Myeloid Leukemia (AML) patients.

## METHODS & IMPLEMENTATION

### Preloaded mutation dataset selection and genomic annotations

Preloaded datasets available in PIO have been downloaded from cBioPortal (http://www.cbioportal.org/) [6,7]. Only diseases with at least one available The Cancer Genome Atlas (TCGA) dataset from the 2018 TCGA pancancer study were selected [8]. The complete list of datasets used and their corresponding studies is provided in the Supplementary Table S1.

Additional genomic information were retrieved from the GRCH37 human genome reference downloaded from the NCBI Reference Sequence Database (Refseq) (GCF_000001405.13). Exon coordinates were extracted using the RefSeq “Select” annotation. Genes with no RefSeq “Select” annotations (such as pseudogenes) were attributed their unique RefSeq ID. Gene and exon length were calculated using the difference between end and start site from each position. Exon ID was attributed using the nearest exon from the mutation start site. Intronic mutations were numbered according to the nearest exon ID and introns were systematically attributed a 500 bp length. Mutations with no feature found at less than 10 kb and with a nearest feature distance exceeding their respective gene’s total size were omitted. All gene names were updated to their last HUGO symbol using the alias2SymbolTable function from the limma R package [9].

### Informativity module algorithm

The PIO algorithm takes as input pre-formatted mutational data from whole-exome or genome sequencing, and it can be run grouping mutations by gene or exon. For the optimal panel selection (optimal mode), the most informative gene on the overall cohort is first selected. Patients presenting this mutation are then removed from the cohort, and the most informative gene in the remaining cohort is then selected. The algorithm reiterates until all patients are removed or no patients are added. The number of unique patients per kilobase (UPKB) or the number of unique patients (UP) can be used as informativity metric. A minimal number of patients per mutation, calculated on the overall cohort, can be set in order to avoid overfitting to private mutations.

For the custom panel interrogation (custom mode), only genes/exons from the provided list of genomic intervals are considered. An additional list of genes/exons, established by running the algorithm on the patients without mutation on the proposed genomic intervals is provided to complete the panel if all patients are not diagnosed with the custom panel.

For the panel test mode, the informativity of the exact list of genes/exons is assessed without further mutation selection.

### Mutation module

The mutation module proposes different statistics calculated on the whole cohort, without running the PIO base algorithm. The main statistics table calculate several statistics such as UP and UPKB on the overall cohort for each gene or exon. Gene or exon annotations are also provided, with the corresponding unique RefSeq exon ID. The cumulated mutations table shows the number of patients with at least x mutation within the considered panel.

### Length analysis module

The length analysis module allows the comparison of four different approaches for panel length optimization. In the classic approaches, mutated genes or exons are ranked according to their informativity metric (either UP or UPKB) computed on the total cohort. Panel informativity is then calculated by taking the number of new unique patients diagnosed for each new gene/exon using the panel tester function. The two other approaches use the PIO algorithm.

### Coding and online tool implementation

The algorithm was written in R language with the R software v.4.0.3 [10]. A while loop was used to perform the recurrent iterations across the input dataset. The graphical user interface (GUI) allowing to use the algorithm on different pre-loaded or custom datasets in a user-friendly environment requiring no coding skill was implemented using the R Shiny package [11]. The resulting application, PIO, was then released as an open-source software under the MIT license and uploaded on github (https://github.com/VincentAlcazer/PIO) and on the shinyapps.io servers (https://vincentalcazer.shinyapps.io/Panel_informativity_optimizer/) to allow both local and online use.

### Pancancer mutation exploration

Pancancer mutations were explored at the gene-level using PIO with default parameters (optimal mode, mutations grouped by gene, UP as informativity metric) on the pre-loaded datasets of 23,203 patients from 91 independent cohorts spanning 31 different cancer types.

### Pancancer optimal panel length comparisons

Panel length to reach full informativity were compared across all cancer types using the PIO length analysis module. For each cancer, an optimal panel was defined using the 4 different approaches proposed by PIO. These approaches were first compared using the overall proportion (%) of patients diagnosed according to panel size in kb smoothed using a general additive model. A focus was then proposed for each metric, comparing the panel size differences allowing to reach 100% informativity between PIO and classical approach. This optimal panel size was then extracted for each cancer type and used for exploratory correlations with the total number of patients and different markers of DNA alterations extracted from Thorsson et al. [12].

### Examples of application

For panel size optimization, a 293-gene (802-exon) panel targeting genes recurrently mutated in hematological malignancies [4] was shrunk using the PIO custom mode with UP or UPKB metric and mutations grouped by exon. The performance of the resulting 140-exon panel was then evaluated in 928 DLBCL patients who benefited from the full panel [3].

For the evaluation of our local routine 110-gene panel, PIO was first used in custom mode (mutations grouped by gene, UP metric, at least 1 patient per mutation) to evaluate the predicted performance in the pre-loaded dataset (predicted). The proposed optimized 33-gene panel was then evaluated in test mode on our local cohort (observed).

For panel benchmarking in AML, 5 commercial myeloid panels [4] together with our local panel were evaluated using PIO in panel test mode (mutations grouped by gene, UP metric, at least 1 patient per mutation) on the preloaded AML datasets in the PIO. Panels where then evaluated using PIO custom mode.

## RESULTS

### PIO: algorithm and software setup

PIO is an open-source software written in R language designed to help in the selection of the most informative set of genes or genomic intervals for the design of NGS panels. A GUI, available at https://vincentalcazer.shinyapps.io/Panel_informativity_optimizer/, provides a user-friendly way to use PIO without any bioinformatics skills (Figure 1). In the main module (“Informativity”), PIO allows to select the most informative genomic intervals (either genes, or exons/introns) from whole-exome or whole-genome sequencing data. Public mutation datasets from 91 independent cohorts and 31 different cancer types are preloaded, and the user can also download additional datasets to explore. The principle of the PIO algorithm is simple. First it selects the most informative genomic interval (*i.e*. with the highest number of mutated patients). After removal of all the patients harboring a mutation in this first genomic interval, the second genomic interval selected is the most informative on the remaining patients. This procedure is reiterated until all patients have at least one mutation selected or if no more patients can be added (see methods). Importantly, the algorithm can be run by gene or by exon/intron, and two informativity metrics can be used: the number of UP or the number of UPKB.

**Figure 1:**
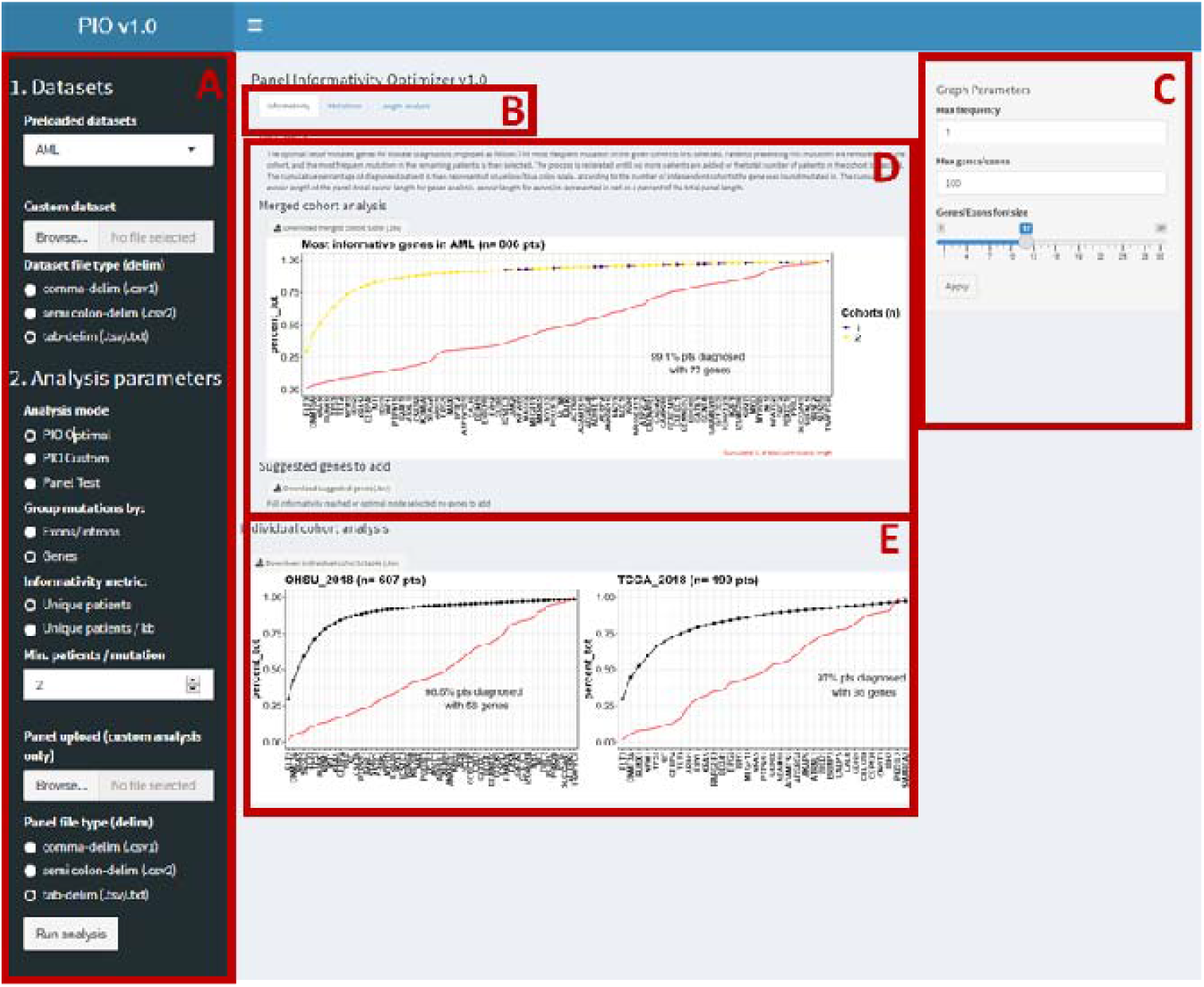
PIO Graphical user interface for dataset interrogation. In panel A, preloaded or custom datasets and panels to be analyzed can be downloaded and the analytical parameters selected. The different analyses proposed by PIO (informativity, mutations, and length analysis) can be selected using tabs (B). Graphical parameters can be tweaked (C). Resulting graphs are shown in the central panels: for the “informativity” tab, the additive effect of the selected genomic intervals on informativity is represented on the overall merged cohort (D) and in each cohort separately (E). A link to download the most informative genomic intervals to add to a custom panel is proposed (D). *NGS: Next Generation Sequencing, PIO: Panel Informativity Optimizer*

The datasets can be interrogated with three different modes: in the so-called “optimal” mode, PIO automatically selects an optimal set of genomic intervals to maximize the informativity in the selected cohort; in the “custom” mode, PIO selects genomic intervals among a custom panel proposed by the user. In the latter case, PIO suggests genomic intervals to add according to their informativity computed on the remaining patients, unless optimal informativity has already been reached. In the panel test mode, the software takes the exact list of genomic intervals from a provided custom panel, ranks them according to the selected metric and presents the panel’s performance in the cohort.

PIO offers several other exploratory analyses: the “mutations” module presents different mutation metrics calculated on the total cohort, such as gene/exon length and UPKB, together with a summary of the cumulated number of mutations. A graphical output illustrates the frequency and nature of the most frequently mutated genomic intervals in the overall cohort, and the data are summarized in a downloadable table. A second graphical output presents a co-occurrence plot of mutations in the selected genomic intervals at the individual level. In addition, the “length” module compares four different approaches to optimize panel design to maximize the informativity/size ratio (see methods).

### Pancancer mutations and optimal panel length exploration

We first used PIO to obtain insight into pancancer mutations at the gene-level. Using PIO default parameters (optimal panel, mutations grouped by gene, UP as informativity metric), we explored the pre-loaded datasets of 23,203 patients from 91 independent cohorts spanning 31 different cancer types (Supplementary Table S1). Analysis of the merged dataset found differences in term of the number of genes required to reach absolute informativity (Figure 2A). Prostate adenocarcinoma (PRAD) and breast invasive carcinoma (BRCA) were the two cancers requiring the largest number of genes to reach exhaustive informativity (284 and 193 genes, respectively). Conversely, uterine carcinosarcoma (UCS) and lung squamous cell carcinoma (LUSC) required a panel of only 5 and 12 genes, respectively, to cover all patients. In 20 cancer types with at least 2 independent cohorts, there was a high level of concordance of the analysis attested by low variance in the number of genes needed to reach exhaustive informativity across cohorts (the median [interquartile range, IQR] standard derivation was 9.66 [5.7-13.2]) (Figure 2B). However, three cancer types showed a more pronounced heterogeneity among cohorts: PRAD, glioblastoma multiforme (GBM), and BRCA (standard derivations: 55.7, 26.2, and 25.5, respectively).

**Figure 2:**
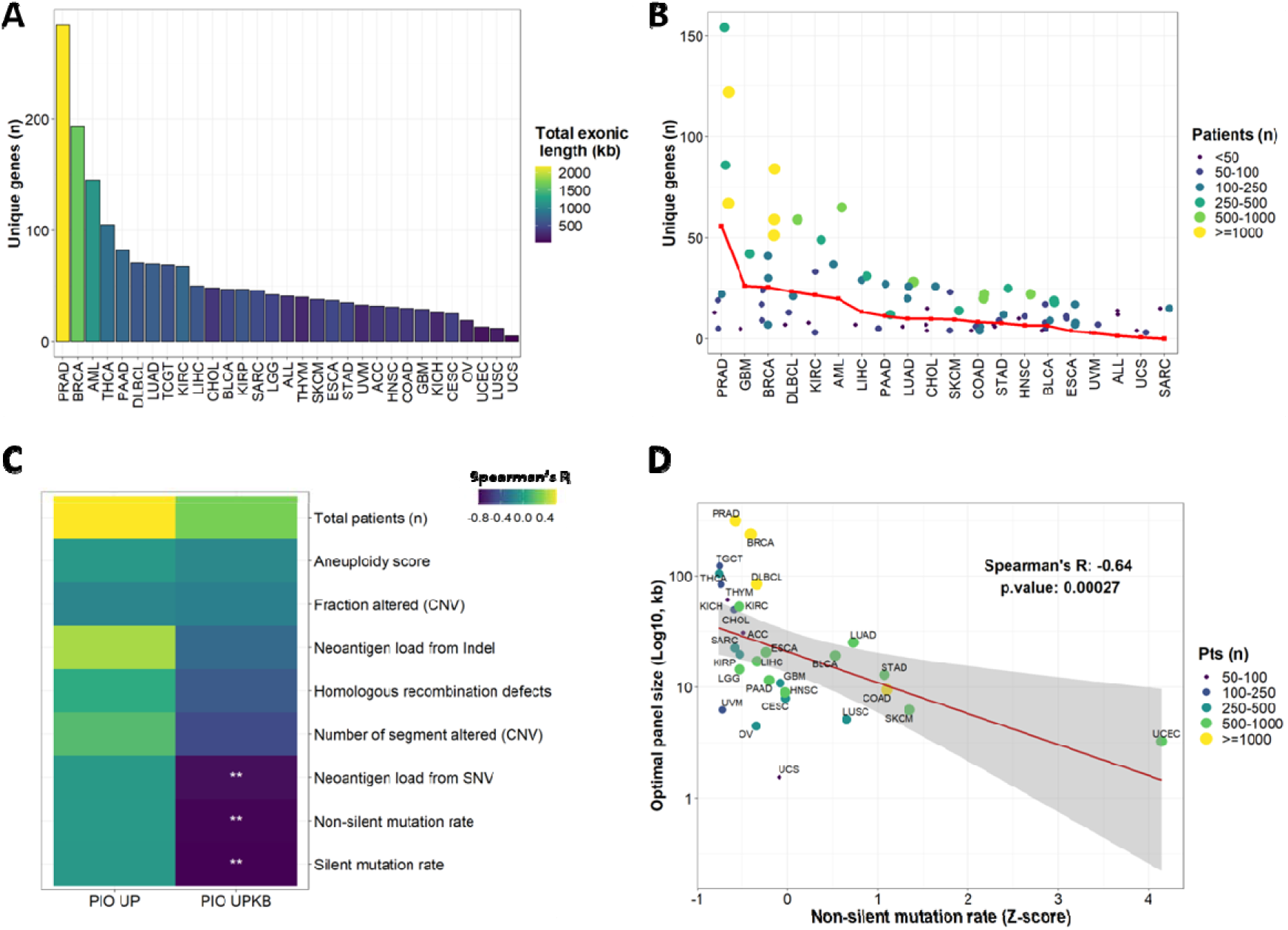
Pancancer exploration at the gene level using PIO. A: Number of unique genes required to reach full informativity in the merged cohorts of specific cancer types. Bars are colored according to the size of the panel. B: Number of unique genes required to reach full informativity in each cohort. Only cancers with at least two independent cohorts are represented (n=20). Standard derivation of gene number is plotted in red for each cancer type. C: Spearman’s correlation between tumor DNA alteration markers (mean value per cancer), total patient number per cohort, and optimal panel length found by PIO using number of unique patients (UP) and number of UP per kilobase (UPKB). Adjusted p-values (False Discovery Rate (FDR)): * < 0.05, ** < 0.01, *** <0.001. D: Spearman’s correlation between the mean non-silent mutation rate and the optimal panel size provided by PIO using the UPKB metric. The linear regression line is shown in red with its 95% confidence interval in grey.

We next used PIO with mutations grouped by exon to select the most informative panel with minimal length for each cancer. The optimal panel length proposed by PIO using the UPKB metric was lower in cancer types with high mutational burden, as assessed by silent/non silent mutation rates or neoantigen load prediction from single nucleotide variants (SNV; Figure 2C and D).

Taken together, these results indicate that PIO allows to explore mutation data and optimize panel design in multiple cancer types.

### Panel length optimization

One of the main goals of PIO is to offer a tool to help optimize the informativity/size ratio. The PIO length analysis module allows the comparison of four panel optimization strategies, with the aim to catch the highest number of patients with the most parsimonious (*i.e*. short) panel. In the “classic” approach, mutated genes or exons are ranked according to their mutation frequency in the total cohort. This strategy is compared with the PIO algorithm (see methods) with an optimization strategy based either on absolute (UP) or relative quantification (UPKB).

We compared these four optimization strategies across all cancer datasets, grouping mutations by exon. As expected, the UPKB informativity metric performed better than UP metric to optimize panel size (Figure 3A). The higher performances of PIO were observed in all types of cancer, allowing panel size reduction up to 995 kb for Ovarian serous cystadenocarcinoma (OV; Figure 3B). The panel size reduction was strongly correlated with the overall panel size (Pearson’s R: −0.93, p=7.1e-14; Figure 3B) and moderately with the total number of patients (Spearman’s R −0.42, p=0.019; Figure 3C). These results show that PIO is an efficient tool to maximize informativity whilst reducing panel length.

**Figure 3:**
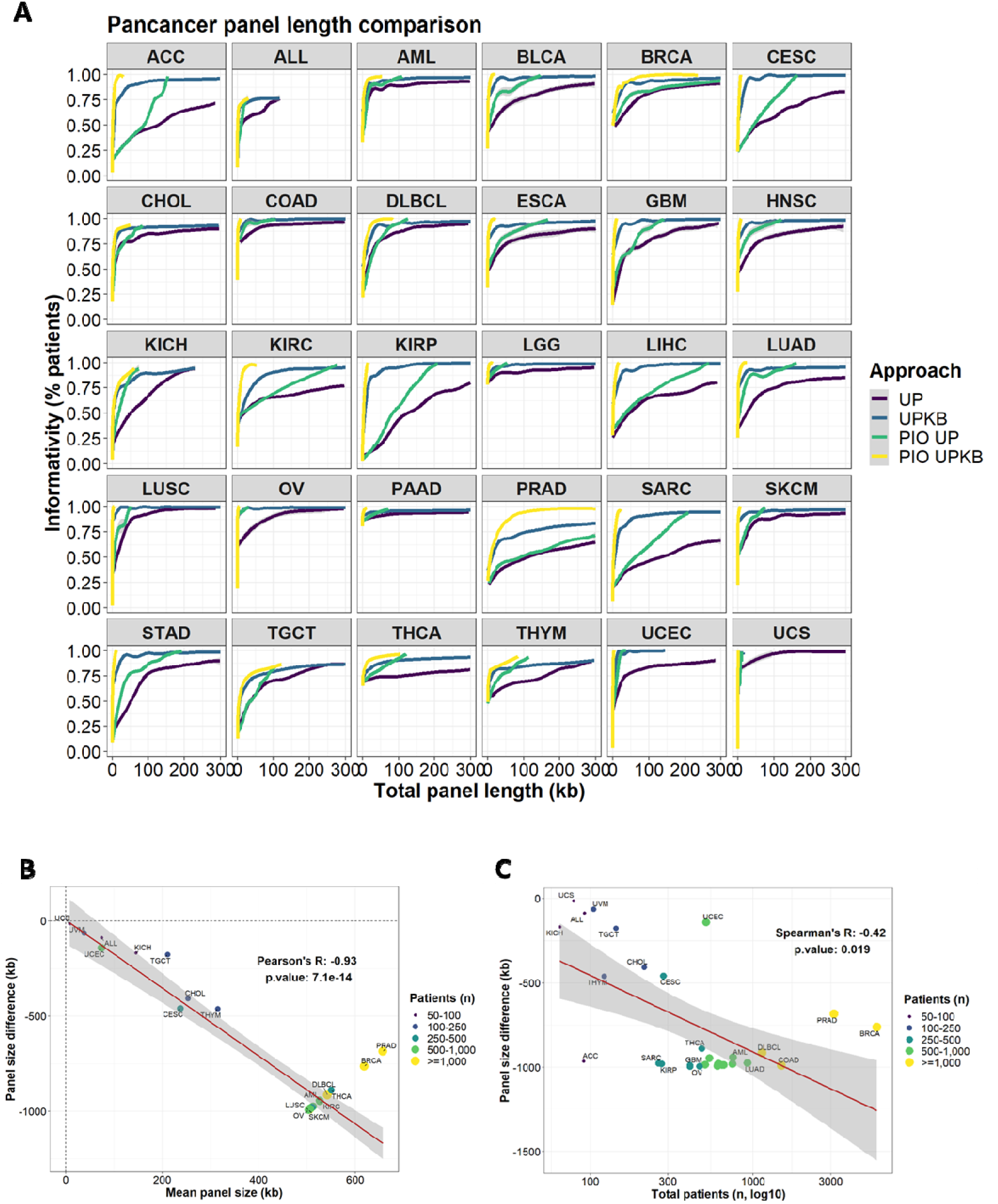
Optimal panel length comparison according to classical and PIO approaches. A: Overall pancancer informativity using the classical or PIO approach, with either number of unique patients (UP) or number of UP per kilobase (UPKB) metric. Overall informativity is represented using a general additive model independently for each individual cancer. B: Differences between PIO and classical approaches for UPKB metric. Each cancer type is plotted according to panel size difference (y-axis) and mean panel size (x-axis). Pearson’s correlation between panel size differences and mean panel size is presented. C: Spearman’s correlation between the total number of patients and panel size difference between PIO and classical approach (using UPKB metrics).

### Examples of application

We assessed the performance of PIO in different settings. We first set out to optimize the size of a 293-gene (802-exon) panel used in DLBCL [3] while retaining maximum informativity. Using the PIO custom mode with UP metric and mutations grouped by exon, we predicted on the preloaded DLBCL datasets that 93.8% of DLBCL patients could be diagnosed with a subset of only 140 exons from this panel (Figure 4A). We compared this prediction to sequencing results of a public dataset of 928 DLBCL who were analyzed by the full panel [3]. In this cohort, 95.8% of patients presented at least one mutation in one of these 140 exons (Figure 4A). Using PIO with either UP or UPKB metric allowed to reduce the panel size from ~300kb to less than 100kb whilst keeping the same level of informativity (Figure 4B).

**Figure 4:**
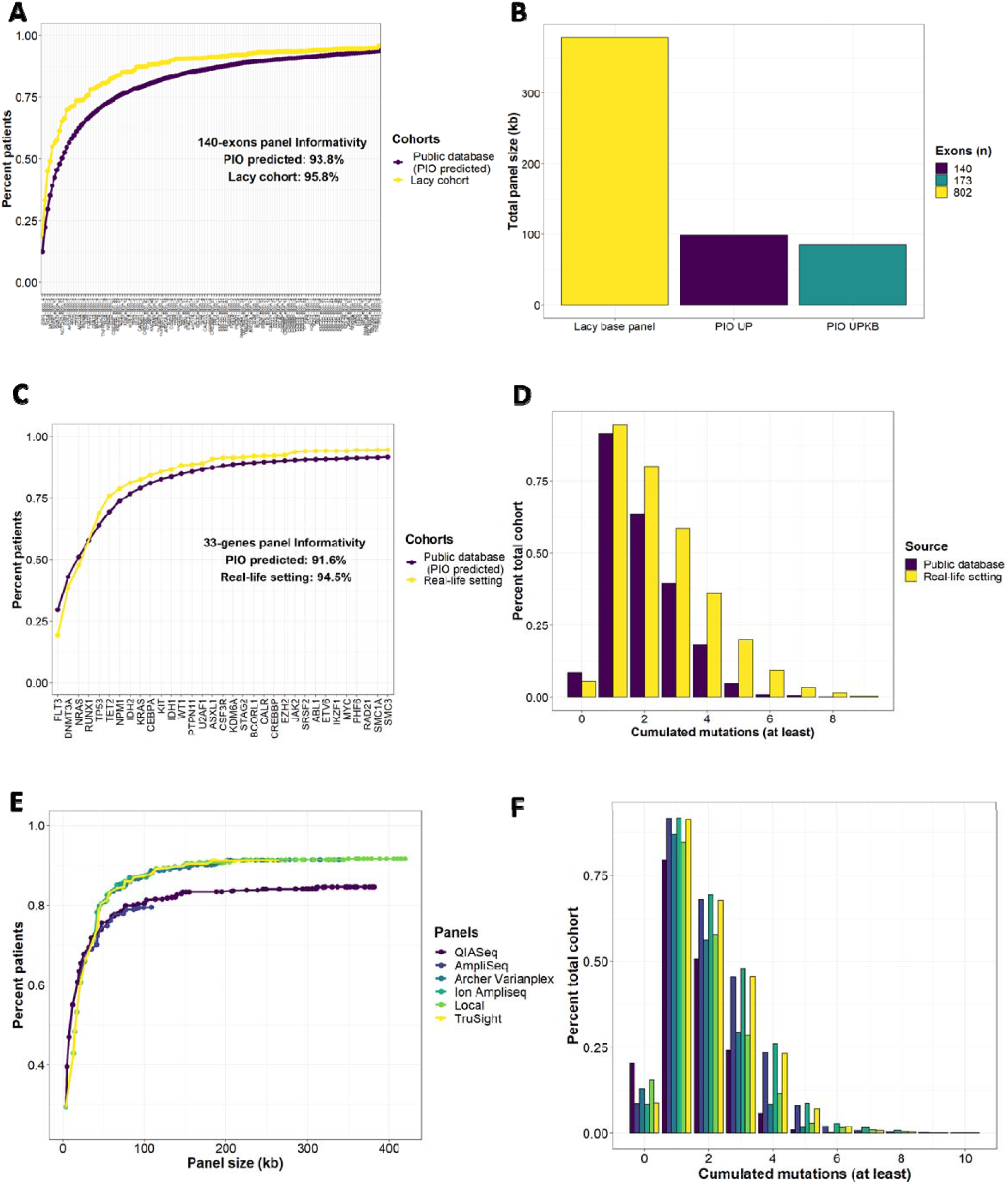
External validation of PIO’s predictions. A: Performance of the 140-exon panel in preloaded Diffuse Large B Cell Lymphoma (DLBCL) datasets and in a published DLBCL cohort (Lacy cohort [3]). The 140-exon panel was selected from the 802-exon panel using the PIO in custom mode (mutations grouped by exon, Unique patients per kilobase (UPKB) as informativity metric, minimum number of patients per mutation set to 1). B: Panel size comparison between the default 802-exon panel used by Lacy et al. [3] and the PIO-optimized panel using either unique patients (UP) (140 exons) or UPKB (173 exons) as metric. C: Performance of the 33-gene panel in the preloaded AML dataset (predicted) and in a local AML cohort (observed). The 33-gene panel was selected from a 110-gene panel using the PIO in custom mode (mutations grouped by gene, UP as informativity metric, minimum number of patients per mutation set to 1). D: Distribution of the number of cumulated mutations in the PIO AML datasets and our local AML cohort using the 33-gene panel. E: Performance of 5 commercial panels and our local panel using PIO on preloaded AML datasets. F: Distribution of the number of cumulated mutations between 5 commercial panels and our local panel in PIO AML datasets.

We then used the PIO in custom mode with UP metric and mutations grouped by gene to evaluate the performance of a custom 110-gene NGS panel used for routine diagnosis of hematological malignancies. Based on the preloaded AML cohorts, PIO predicted that 91.6% patients would have at least one mutation with only 33 of these genes (Figure 4C). Indeed, 700/714 (94.5%) AML patients referred to our laboratory between 2014 and 2020 had at least one mutation in one of these 33 genes (Figure 4C). The total number of mutations per patient was also roughly similar between PIO predictions and the local cohort data for up to 3 cumulated mutations per patient (Figure 4D).

Finally, we used the PIO panel test mode to compare 5 commercial myeloid panels [4] together with our custom panel to the preloaded AML datasets. There was an overall better informativity for Trusight, Ion AmpliSeq, Archer Varianplex and our custom panel, though no panel reached full informativity (Figure 4E). Differences in terms of cumulated number of mutations were also observed (Figure 4F). By runing PIO in custom mode with each panel on the preloaded AML datasets, we observed that optimal informativity could be reached with only half of each panel size (Supplementary Figure S1).

## DISCUSSION

We propose herein PIO, an open-source software to help establishing NGS panels and which offers several applications such as mutation exploration, panel length optimization, and commercial panel benchmarking.

Newman et al. proposed a method to optimize panel informativity in the design phase of CAPP-Seq with an approach that aims to maximize the number of mutations per patient in order to increase the sensitivity of the analysis of ctDNA for minimal residual disease monitoring [5]; this differs to PIO that aims to maximize the informativity/panel size ratio. Accordingly, PIO will select smaller panel sizes than the one proposed by CAPP-seq, but at the cost of identifying fewer targets for minimal residual disease monitoring. Moreover, PIO stands out by proposing an open-source tool with a user-friendly interface. As shown in the present study, PIO can also be used to explore mutation frequencies and other metrics in a given cancer or across a pancancer dataset. Furthermore, as an open-source software, any user may integrate the code on a routine pipeline, suggest improvement or new features implementation online or directly modify/adapt the code to match its specific needs. Compared to an empirical approach based on raw UP or UPKB metrics, PIO systematically improved panel size while keeping the same informativity. We observed a correlation between the magnitude of panel size reduction and the size of the cohort which can be explained by higher heterogeneity in large cohorts, necessitating an important number of genes to reach an acceptable informativity. The use of PIO is of particular interest for optimizing panel size in these cancer types.

Considering potential cohort bias is also important when using PIO. Although a low variance in the number of genes needed for full informativity was observed for most cancers, some were characterized by high heterogeneity between cohorts. This heterogeneity could be explained by disease heterogeneity (several molecular subtypes) or differences in patient inclusion criteria. To circumvent this, PIO allows to interrogate any custom dataset, hence enabling the user to select representative studies according to its specific clinical settings.

While allowing to maximize informativity, PIO does not account for specific settings such as the prognosis or theragnostic value of a given mutation. Hence, although clinically relevant events will be included according to their relative frequency and informativity, the selection of a set of gene by PIO does not preclude the manual inspection of the panel by a trained molecular biologist to verify and add clinically relevant and mandatory mutations.

## CONCLUSION

PIO is an efficient and accessible free open-source software that allows the optimization of cancer NGS panel informativity and size, explore mutation data, and benchmark different panels for a given cancer. As no other broadly accessible software exists, PIO should be of major interest for molecular biologists and physicians to help in routine cancer NGS diagnosis panel design.

## AVAILABILITY AND REQUIREMENTS

Project name: Panel Informativity Optimizer

Project home page: https://github.com/VincentAlcazer/PIO

Operating system(s): Platform independent

Programming language: R

Other requirements: NA

License: MIT

Any restrictions to use by non-academics: No

## LIST OF ABBREVIATIONS

AML: Acute Myeloid Leukemia
BRCA: Breast invasive carcinoma
DLBCL: Diffuse Large B-Cell Lymphoma
GBM: Glioblastoma Multiforme
IQR: Interquartile Range
LUSC: Lung squamous cell carcinoma
NGS: Next generation sequencing
PIO: Panel informativity optimizer
PRAD: Prostate adenocarcinoma
Refseq: Reference sequence database
TCGA: The cancer genome atlas
SNV: Single nucleotide variant
UCS: Uterine carcinosarcoma
UP: Unique patients
UPKB: Unique patients per kilobase

## DECLARATIONS

### Ethics approval and consent to participate

Not applicable

### Consent for publication

Not applicable

### Availability of data and materials

The dataset(s) supporting the conclusions of this article together with software code are available in the Github repository https://github.com/VincentAlcazer/PIO.

### Authors’ contributions

**Vincent Alcazer:** Methodology, Software, Validation, Formal analysis, Investigation, Resources, Data curation, Writing - Original Draft, Visualization; **Pierre Sujobert** Conceptualization, Methodology, Validation, Writing - Review & Editing, Supervision, Project administration

## Acknowledgements

The results published here are in part based upon data generated by the TCGA Research Network: https://www.cancer.gov/tcga. The authors would like to thanks Philip Robinson for his help with English editing.

## Supplementary material

**Additional file 1: supplementary figures (.pdf)**

**Additional file 2: supplementary tables (.docx)**

**Supplementary figure S1: Commercial panel optimization**

A: Performance of 5 commercial panels and our local panel using PIO in optimal mode (mutations groups by gene, UP as informativity metric, minimum number of patients per mutation set to 1).

**Supplementary table S1: Complete list of preloaded datasets included in the PIO**

