## Supplementary Table S1 for "Panel Informativity Optimizer (PIO): an R package to improve cancer NGS panel informativity"

| **Abbreviation** | **Full cancer name** | **Cohorts** | **Total patients** | **References** |
| --- | --- | --- | --- | --- |
| ACC | Adrenocortical carcinoma | 1 | 91 | [1] |
| ALL | Acute Lymphoblastic Leukemia | 2 | 110 | [2,3] |
| AML | Acute Myeloid Leukemia | 2 | 805 | [4,5] |
| BLCA | Bladder Urothelial Carcinoma | 7 | 765 | [6–11] |
| BRCA | Breast invasive carcinoma | 9 | 5829 | [12–20] |
| CESC | Cervical squamous cell carcinoma and endocervical adenocarcinoma | 1 | 281 | [1] |
| CHOL | Cholangiocarcinoma | 4 | 241 | [1,21–23] |
| COAD | Colon adenocarcinoma | 6 | 1492 | [24–29] |
| DLBCL | Diffuse Large B-cell Lymphoma | 4 | 1188 | [1,30–33] |
| ESCA | Esophageal carcinoma | 5 | 615 | [34–38] |
| GBM | Glioblastoma multiforme | 2 | 410 | [39,40] |
| HNSC | Head and Neck squamous cell carcinoma | 3 | 608 | [41–43] |
| KICH | Kidney Chromophobe | 1 | 65 | [1] |
| KIRCH | Kidney renal clear cell carcinoma | 4 | 550 | [44–47] |
| KIRP | Kidney renal papillary cell carcinoma | 1 | 274 | [1] |
| LGG | Brain Lower Grade Glioma | 1 | 510 | [1] |
| LIHC | Liver hepatocellular carcinoma | 3 | 610 | [1,48,49] |
| LUAD | Lung adenocarcinoma | 4 | 942 | [50–53] |
| LUSC | Lung squamous cell carcinoma | 1 | 469 | [1] |
| OV | Ovarian serous cystadenocarcinoma | 1 | 409 | [54] |
| PAAD | Pancreatic adenocarcinoma | 3 | 658 | [1,55,56] |
| PRAD | Prostate adenocarcinoma | 8 | 3289 | [57–64] |
| SARC | Sarcoma | 2 | 269 | [1,65] |
| SKCM | Skin Cutaneous Melanoma | 3 | 609 | [1,66,67] |
| STAD | Stomach adenocarcinoma | 5 | 663 | [68–72] |
| TGCT | Testicular Germ Cell Tumors | 1 | 144 | [1] |
| THCA | Thyroid carcinoma | 1 | 485 | [1] |
| THYM | Thymoma | 1 | 123 | [1] |
| UCS | Uterine Carcinosarcoma | 2 | 79 | [1,73] |
| UCEC | Uterine Corpus Endometrial Carcinoma | 1 | 515 | [74] |
| UVM | Uveal Melanoma | 2 | 105 | [1,75] |

**Supplementary table S1 :** **Complete list of datasets included in PIO**
