## Supplementary figures and images for "Panel Informativity Optimizer (PIO): an R package to improve cancer NGS panel informativity"

### Supplementary Figure S1

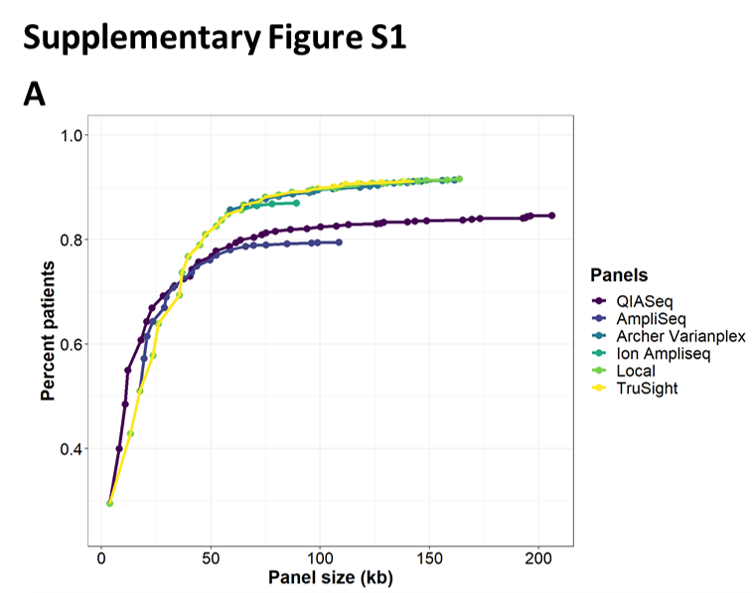
